# Antimicrobial mechanism of *in-situ* plasma activated water treatment of pathogenic *Escherichia coli* and *Staphylococcus aureus* biofilms

**DOI:** 10.1101/2024.07.07.602420

**Authors:** Binbin Xia, Heema Kumari Nilesh Vyas, Scott A. Rice, Timothy P. Newsome, Patrick J. Cullen, Anne Mai-Prochnow

**Author notes:** Corresponding author: E-mail address (B. Xia), (A. M. Prochnow).

## Abstract

**Aims:** This study investigated the efficacy and mechanisms of inactivation of against *Escherichia coli* UTI89 and *Staphylococcus aureus* NCTC8325 through an *in-situ* plasma-activated water (PAW) treatment.

**Methods and Results:** PAW was prepared by discharging atmospheric pressure cold plasma beneath the surface of sterile distilled water. The study investigated the inactivation of biofilm cells and biofilm matrix. A complete killing of biofilm cells was achieved on both of *E. coli* (6.76 ± 0.01 log CFU/mL) and *S. aureus* (6.82 ± 0.02 log CFU/mL). This process happened earlier in *S. aureus*. Simultaneously, PAW treatment disrupted the biofilm structure, inducing a significant reduction in general biofilm biomass and extracellular polymer substances (EPS) matrix. With the disruption of EPS, PAW was enabled to further interact with the bacterial membrane, causing a significant increase in membrane permeability and disrupted membrane structure. Finally, PAW treatment led to a significant accumulation of intracellular reactive oxygen and nitrogen species within the biofilm cells.

**Conclusions:** Collectively, these findings indicate that PAW effectively inactivates biofilms by mechanically targeting the biofilm EPS matrix and biofilm cells in both gram-negative and gram-positive bacteria.

**Impact statement:** This study contributes novel insights into plasma-activated water’s mechanisms of action, particularly its impact on the biofilm extracellular polymeric substances matrix (exopolysaccharides, extracellular DNA, and protein), cell membrane permeability, depolarization, and intracellular ROS and RNS accumulation in both of Gram-positive and Gram-negative species. These findings highlight PAW-based treatments against biofilm-related challenges in antimicrobial development and water system decontamination.

## 1 Introduction

Pathogenic biofilms pose a significant challenge in healthcare settings and various industries. They are the main cause of the rise in microbial contamination forming in water distribution systems, including pipes, tanks, and filters. Drinking contaminated water or engaging in water activities like swimming, sailing, and other water sports in contaminated waters can result in waterborne diseases caused by bacteria and other pathogenic microorganisms (Carrascosa 2021; Dula 2021; Shineh 2023). Unlike planktonic cells, bacterial biofilms, form structured communities with a protective matrix, rendering them 1000-fold more resistant to conventional cleaning and disinfection methods (Billings 2015; Cámara 2022). The protective matrix of the biofilm limits the penetration of disinfectants, and bacterial cells within the biofilm may enter a dormant state, making them less susceptible to chemical treatments.

At the core of biofilm architecture lies the extracellular polymeric substance (EPS). This matrix, secreted by microorganisms themselves or captured from host components, plays a crucial role in shaping the behaviour, resilience, and functionality of biofilms. The EPS matrix can be comprised of polysaccharides, DNA, RNA, proteins, and lipids, which affect the permeability and mechanical properties of the biofilm (Billings 2015; Karygianni 2020; Limoli 2015). All of these polymers act as the adhesive that binds microbial cells together, providing structural integrity to the biofilm community (Danese 2000; Horvat 2019). Most antimicrobial agents kill cells, but the matrix may be left behind and that may help with biofilm regrowth and recruitment of new cells (Campoccia 2021). Therefore, it is important to find a treatment that will get rid of the matrix as well as kill cells.

Innovative approaches are continuously sought to combat biofilm-related issues. One promising solution that has garnered attention is plasma-activated water (PAW) (Chen 2018; Guo 2021a). Plasma, often referred to as the fourth state of matter, is a highly energetic state of ionized gases, including excited and neutral atoms, electrons, free radicals, ions, and high-energy photons (Jenns 2022; Muhammad 2018). When applied to water, plasma generates a reactive mixture rich in oxidizing species and free radicals. These physiochemical reactions occur between gaseous substances and liquid molecules and during the transfer of gaseous substances to liquids, resulting in a higher density of reactive oxygen and nitrogen species (RONS) of PAW (Mai-Prochnow 2021; Zhou 2020). This unique composition enables PAW to exhibit potent antimicrobial properties, providing a potential avenue for disrupting and eradicating biofilms.

Despite the well documented bactericidal effect of PAW against biofilms of several species, including *Escherichia coli, Pseudomonas aeruginosa, Listeria monocytogenes, and Staphylococcus aureus* (Guo 2021b; Handorf 2021; Vyas 2023; Ziuzina 2015), there remains a notable gap in understanding the mechanism of PAW’s antibacterial effect on biofilms. Current research suggests that PAW mechanically induces an etching effect to prompt the detachment of biofilms from surfaces. Subsequently, RONS present in PAW, (e.g., peroxide, nitrate, OH radicals) are free to further penetrate the EPS matrix and disrupt the cell membrane integrity (Mai-Prochnow 2021)]. The activity of PAW varies depending on factors such as the type of plasma generated, bacterial strains, and the biofilm EPS matrix components. At this stage, it remains unclear why the effects of PAW differ for these factors.

Our objective was to investigate the efficacy and mechanisms of *in situ* PAW activity against *E. coli* UTI89 and *S. aureus* NCTC8325 biofilm. We demonstrate that PAW initially eliminates a significant portion of *E. coli* and *S. aureus* biofilm cells, resulting in disruption of biofilm structure, specifically EPS matrix. By proving its efficacy against both Gram-positive and Gram-negative bacterial biofilms and exploring its mechanisms of action, our results highlight the potential of PAW in biofilm control strategies and alleviate the associated challenges in healthcare, food safety, and water treatment.

## 2 Materials and Methods

### 2.1 Bacterial strains and cultivation

Uropathogenic *Escherichia coli* UTI89 and pathogenic *Staphylococcus aureus* NCTC8325 were chosen as the Gram-negative and Gram-positive strains, respectively. *E. coli* and *S. aureus* were routinely cultured on Luria-Bertani (LB) agar, consisting of 10.0 g/L pancreatic digest of casein tryptone, 10.0 g/L sodium chloride, 5.0 g/L yeast extract, and 7.5 g/L agar powder. A single colony from a freshly streaked agar plate was inoculated into 5 mL of LB broth and incubated overnight at 37°C with agitation at 160 rpm.

### 2.2 Biofilm formation

*E. coli* and *S. aureus* biofilms were formed on sterile stainless-steel coupons (Thickness = 3.8 mm, Diameter = 12.7mm; BioSurface Technologies, USA) placed in a 24-well plate. 1 mL of the diluted culture (∼ 5×10^6^ CFU/mL) was inoculated into each well of the plate and then incubated for 48 hours at 37 °C without shaking to facilitate cell attachment and subsequent biofilm formation.

### 2.3 PAW generation and treatment

PAW was produced using a bubble spark discharge (BSD) reactor, following the procedure outlined in our prior research (Xia 2023). Briefly, the BSD reactor was immersed in 100 mL of sterile MilliQ water contained within a 250 mL Schott bottle, with coupons bearing attached biofilms (Figure 1). A high-voltage power source (Leap100, PlasmaLeap Technologies, Sydney) was utilized to discharge plasma into the water, employing an input voltage of 150 V, a discharge frequency of 1500 Hz, a resonance frequency of 60 kHz, and a duty cycle of 100 µs. Treatment durations ranged from 1 to 20 min with a compressed air gas flow of 1 standard liter per minute (slm). Coupons with biofilms were immersed in 100 mL of sterile MilliQ water and subjected to plasma treatment. As a control, coupons were submerged in 100 mL of sterile MilliQ water without plasma discharge.

**Figure 1.**
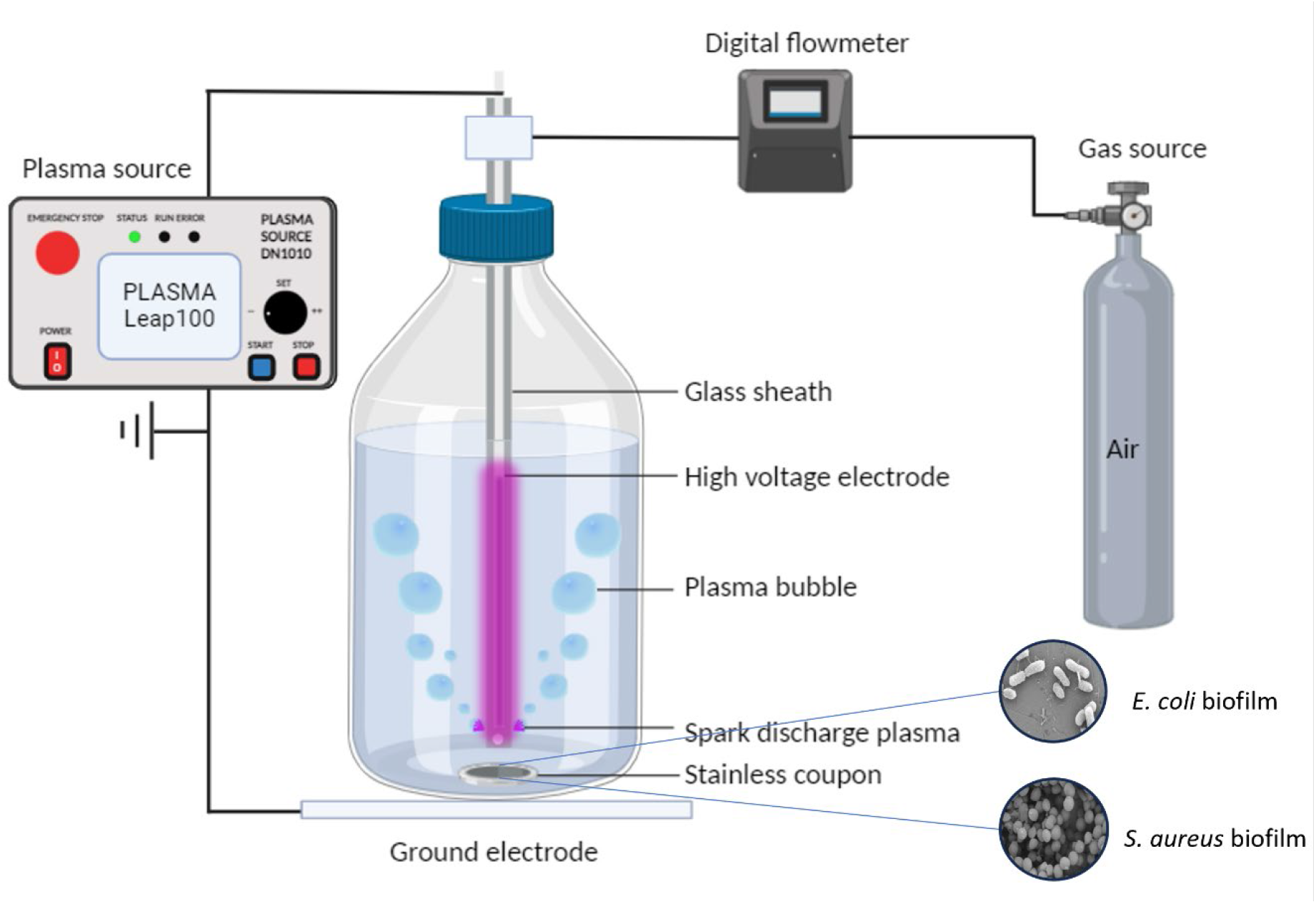
A schematic illustration of the *in-situ* biofilm treatment using plasma-activated water (PAW) generated by a bubble spark discharge (BSD) plasma reactor. *E. coli* UTI89 biofilm and *S. aureus* NCTC8325 biofilm were formed on stainless-steel coupons and positioned at the bottom of a Schott bottle containing 100 mL of MilliQ water. The BSD reactor comprises a high-voltage electrode and a glass sheath. To induce spark discharge plasma, a flow rate of 1 standard liter per minute (slm) of compressed air was utilized to bubble the water, while the high-voltage electrode was energized by the PlasmaLeap100 power source.

### 2.4 Biofilm cell viability

Immediately following the PAW treatment, coupons were retrieved from the treatment bottle and transferred into a Falcon tube containing 1 mL of 1x phosphate-buffered saline (PBS). The biofilm was detached from the coupon surface by gently scraping it with a sterile flat-end spatula. Subsequently, a 3 min water bath sonication at 45 kHz, followed by 10 s of vortexing, was employed to ensure complete dislodgment of biofilms. It is noteworthy that this procedure did not affect cell viability (data not shown). Serial dilutions were then drop-plated (10 µL) onto LB agar in triplicates. The plates were subsequently incubated overnight at 37 °C before determining the colony-forming units (CFU). The CFU log10 reduction (Equation 1) and percentage of reduction (Equation 2) were calculated as follows:

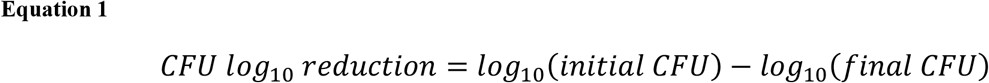

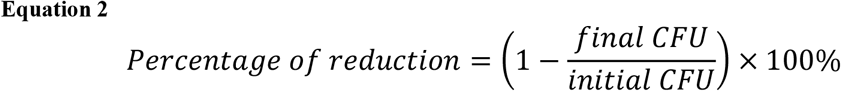

### 2.5 Biofilm biomass crystal violet assay

Following PAW treatment, coupons were retrieved from the treatment bottle and transferred into a Falcon tube containing 1 mL of 1× PBS. The biofilms were then air-dried thoroughly at room temperature for 30-40 min or until completely dried. Subsequently, the air-dried biofilms were stained with 900 µL of 0.2% crystal violet (v/v, Sigma-Aldrich, USA) dissolved in distilled water supplemented with 1.9% ethanol (v/v, Sigma-Aldrich) for 10 min at room temperature under static conditions. After staining, excess crystal violet was removed, and the biofilms were gently washed three times with 900 µL of PBS. The crystal violet stain incorporated into the biofilm was re-solubilized by adding 900 µL of 1% sodium dodecyl sulfate (SDS) (w/v, Sigma-Aldrich, USA), followed by a 10 min incubation at room temperature in the dark. Biofilm biomass was quantified by spectrophotometrically measuring the re-solubilized crystal violet stain, which had previously incorporated into the sample, at OD540 nm/570 nm using a CLARIOStar microplate reader. The biomass measurement encompasses both bacterial cells and the EPS matrix.

### 2.6 Biofilm EPS matrix

#### 2.6.1 EPS matrix

EPS matrix was quantitated by staining with 5 µg/mL Concanavalin A (Con A) Conjugated Alexa Fluor 647 (C21421, Invitrogen, USA) for 15 min in the dark, following adaptations from (Costa Oliveira 2017). The EPS matrix was measured at Excitation 650 nm/ Emission 668 nm (CLARIOStar microplate reader). The EPS matrix was calculated as the mean fluorescent intensity of stained biofilms subtract the mean background intensity of unstained biofilms.

#### 2.6.2 Confocal laser scanning microscopy

To qualitatively visualize the distribution of biofilm extracellular polymeric substance (EPS) components, including extracellular DNA (eDNA), protein, and polysaccharides, additional fluorescent probes were utilized: Sytox Blue (S11348, Invitrogen, USA), Sypro Ruby (F10318, Invitrogen, USA), and Concanavalin A (Con A) Alexa Fluor 647, respectively (Costa Oliveira 2017; Vyas 2020).

Briefly, prior to staining, the PAW-treated biofilms were washed three times with 0.5 mL of PBS. EPS matrix of eDNA and protein were stained with 1 mL of 5 µM Sytox Blue and 1x concentration of Sypro Ruby stain, respectively, for 30 minutes in the dark. Meanwhile, the polysaccharide component of the matrix was stained with 5 µg/mL Con A Alexa Fluor 647 for 15 minutes in the dark. Coupons without biofilm served as controls for background staining. Upon completion of the staining process, the stained biofilms were visualized using an inverted Nikon scanning laser microscope. Specifically, Sytox Blue was imaged at Ex/Em 440nm/482nm, SYPRO Ruby at Ex/Em 450nm/610nm, and Con A Alexa Fluor 647 at Ex/Em 625nm/680nm. Random positions of each sample were imaged.

### 2.7 Scanning electron microscopy

Scanning electron microscopy (SEM) imaging was performed to evaluate the morphological and architectural alterations induced in *E. coli* and *S. aureus* biofilms following 1 and 5 minutes of *in situ* PAW treatment, with comparisons made to a MilliQ control treatment. Briefly, as modified from (Vyas 2023), 48 h biofilms air dried and pre-fixed for 30 min at 4 °C, and then fixed for 1 h at 4 °C. Post-fixation, washed biofilms were dehydrated via graded ethanol series (25%, 50%, 75%, and 3 × 100%). Critical point dried biofilm samples were then sputter coated with 10 nm of gold using a CCU-010 HV high compact vacuum coating system (Safematic, Switzerland). Samples were imaged using the Zeiss Sigma VP HD scanning electron microscope (ZEISS, Germany) at 500 x and 15000 x magnification. Images were captured at random positions to minimize bias.

### 2.8 Membrane permeability study

The outer membrane permeability of *E. coli* and *S. aureus* was assessed using the N-phenyl-1-naphthylamine (NPN) uptake assay following the method described by Halder (2015). Both *E. coli* and *S. aureus* were cultured to logarithmic phase, reaching approximately 10^8^ cells/mL. Subsequently, 100 μL of the bacterial suspension (≈5 × 10^6^ CFU/mL) was dispensed into each well of a 96-well plate. The plate was then centrifuged at 3900 g for 15 minutes, and the supernatant from each well was carefully removed and replaced with 50 µL of 10 µM NPN solution (dissolved in PBS). The stained cells were subsequently exposed to 200 µL of PAW and control Bubble water, with fluorescence measurements taken at various time points (0, 30, 60, 120, and 180 min). As a positive control, 20 µg/mL of cetyltrimethylammonium bromide (CTAB) was used (Halder 2015). The change in NPN fluorescence was quantified at excitation/emission wavelengths of 350-15 nm/420-15 nm. All experiments were performed in triplicate for statistical analysis. Membrane permeability was determined by calculating the difference between the fluorescence intensity of NPN-stained cells and the background fluorescence intensity of the cell suspension without NPN.

Membrane depolarization in *E. coli* and *S. aureus* was evaluated using 3,3’-diethylthiadicarbocyanine iodides (2 μmol/L; Sigma), a fluorogenic dye capable of measuring changes in transmembrane potential as previously described (Vyas 2023). The dye was allowed to incorporate into bacteria cells for 20 minutes at 37°C. Subsequently, the bacteria were washed and exposed to PAW, as well as control treatments (MilliQ and Bubble). Fluorescence intensity was measured at excitation/emission wavelengths of 600-15/660-15 nm using a ClarioStar microplate reader, and membrane depolarization was quantified in arbitrary units. As a positive control, 200 µg/mL of cetyltrimethylammonium bromide (CTAB) was utilized (Halder 2015). Membrane depolarization was determined by subtracting the fluorescence intensity of stained cells from the background fluorescence intensity of the stainless cell suspension.

### 2.9 Intracellular ROS and RNS detection

The accumulation of intracellular ROS and RNS PAW-treated *E. coli* and *S. aureus* biofilm cells were investigated using 2′,7′–dichlorofluorescin diacetate (DCFDA; Sigma Aldrich, Australia) and 4,5-diaminofluorescein diacetate (DAF-FM; Sigma-Aldrich, Australia) staining assays, as previously described (Vyas 2023). In brief, 48 h *E. coli* and *S. aureus* biofilms were exposed to 200 µL of PAW or MilliQ water (MQ) as a control for 15 minutes. Following PAW exposure, the biofilms were stained with 150 µL of 20 µM DCFDA or 5 µM DAF-FM for 30 minutes. The levels of intracellular ROS and RNS were quantified using a microplate reader at Ex/Em of 485-15 nm/545-15 nm and Ex/Em 495-15 nm/515-15 nm, respectively. The final fluorescent intensity was determined by subtracting the background intensity from the fluorescent intensity of the stained cell solution.

### 2.10 Statistical analysis

Experiments were performed 3 times and values are expressed as mean ± standard error of mean 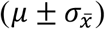. A parametric, One-way ANOVA (with Tukeys multiple comparisons test, p<0.05) was performed where appropriate to identify significant differences in log reduction of each sample compared to the control. Two-way ANOVA (with Tukeys multiple comparison test, p < 0.05) was performed where appropriate to identify significant differences in bacterial membrane permeability and intracellular ROS and RNS level of each sample.

## 3 Results

### 3.1 *In situ* PAW treatment inactivated biofilm cells

The efficacy of PAW in inactivating biofilms of *E. coli* UTI89 and *S. aureus* NCTC 8325 bacterial strains was evaluated. A time-dependent inactivation of biofilm cells was observed upon *in situ* PAW treatment. A complete reduction of CFU was achieved after 13 min of treatment for *E. coli* (6.76 ± 0.01 log CFU/mL) and after 11 min treatment for *S. aureus* biofilms (6.82 ± 0.02 log CFU/mL) (Figure 2). PAW treatment for 5 minutes already resulted in a significant log reduction in both *E. coli* (3.06 ± 0.06 log CFU/mL) and *S. aureus* (3.78±0.03 log CFU/mL), corresponding to over 99% reduction compared to the initial CFU. The biofilm populations of *E. coli* exhibited greater susceptibility than *S. aureus* biofilms within the first minute of PAW treatment. However, S. *aureus* biofilm cells were reduced to below the detection limit of 1.0 log10 at 11 min, while it took 13 min for *E. coli* biofilms to reach the same level. No significant difference was observed in the reduction of overall biofilm populations between these two bacterial species.

**Figure 2.**
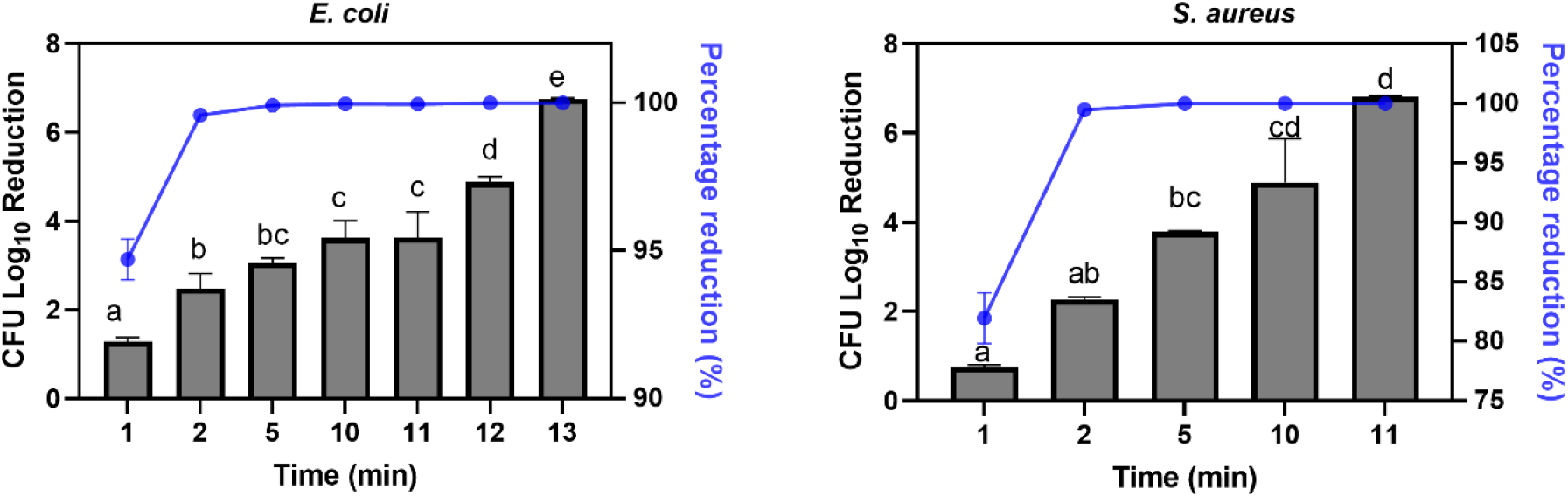
Inactivation efficacy of PAW with different plasma discharge time on 48h *E. coli UTI89* biofilm and *S. aureus NCTC8325* biofilms. Different letters indicate significant differences between groups (p<0.05). The data are represented as mean ± SEM (n=3).

### 3.2 Effect of PAW on biofilm biomass

The total biofilm biomass is another critical factor in studying inactivation efficiency. Biofilm biomass was assessed using the crystal violet staining assay. As shown in Figure 3, biofilms formed by both *E. coli* and *S. aureus* were reduced significantly after PAW treatment. More specifically, the biomass of *E. coli* biofilms decreased rapidly by 48.4% after 1 min treatment and reduced by up to 53.5% after 15 min treatment compared to the control (0 min treatment). Similarly, PAW-treated *S. aureus* biofilm biomass exhibited a time-dependent reduction, with reductions of 23.6% after 1 min treatment and 50.2% after 15 min treatment compared to the control (0 min treatment). However, after the initial decrease in biomass detected by CV, no further reduction was observed and the absorbance numbers for the 15 min treatment are not significantly different to the 5 min treatment.

**Figure 3.**
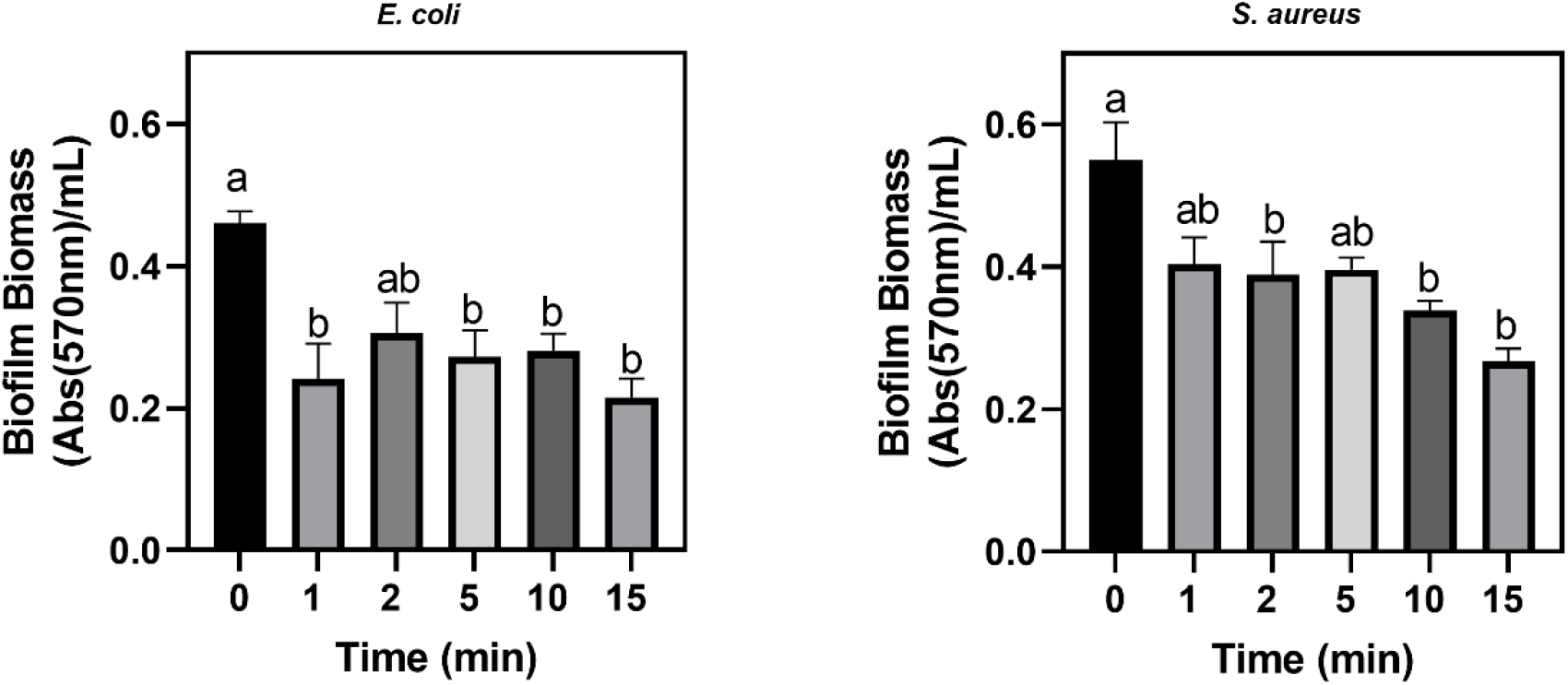
Change in biofilm biomass of *E. coli* UTI89 and *S. aureus* NCTC8325 biofilm challenged by 0 to 15 min of *in situ* PAW treatment. Data represents mean ± SEM. Different letters indicate significant differences between groups (p<0.05); n =2 biological replicates, with 2 technical replicates each.

### 3.3 EPS matrix response to PAW treatment

To better understand the effect of PAW on the biofilm, the EPS matrix was assessed using a fluoroprobe, Con A Alexa Fluor 647, known for its reactivity with aldehydes in polysaccharides. As shown in Figure 4, PAW treatment notably reduced the EPS matrix in both *E. coli* and *S. aureus* biofilms in a time-dependent manner., significant differences in polysaccharides response to PAW treatment were observed and the EPS was reduced by 80.69% for E. coli and 79.08% for S. aureus after 15 min PAW treatment.

**Figure 4.**
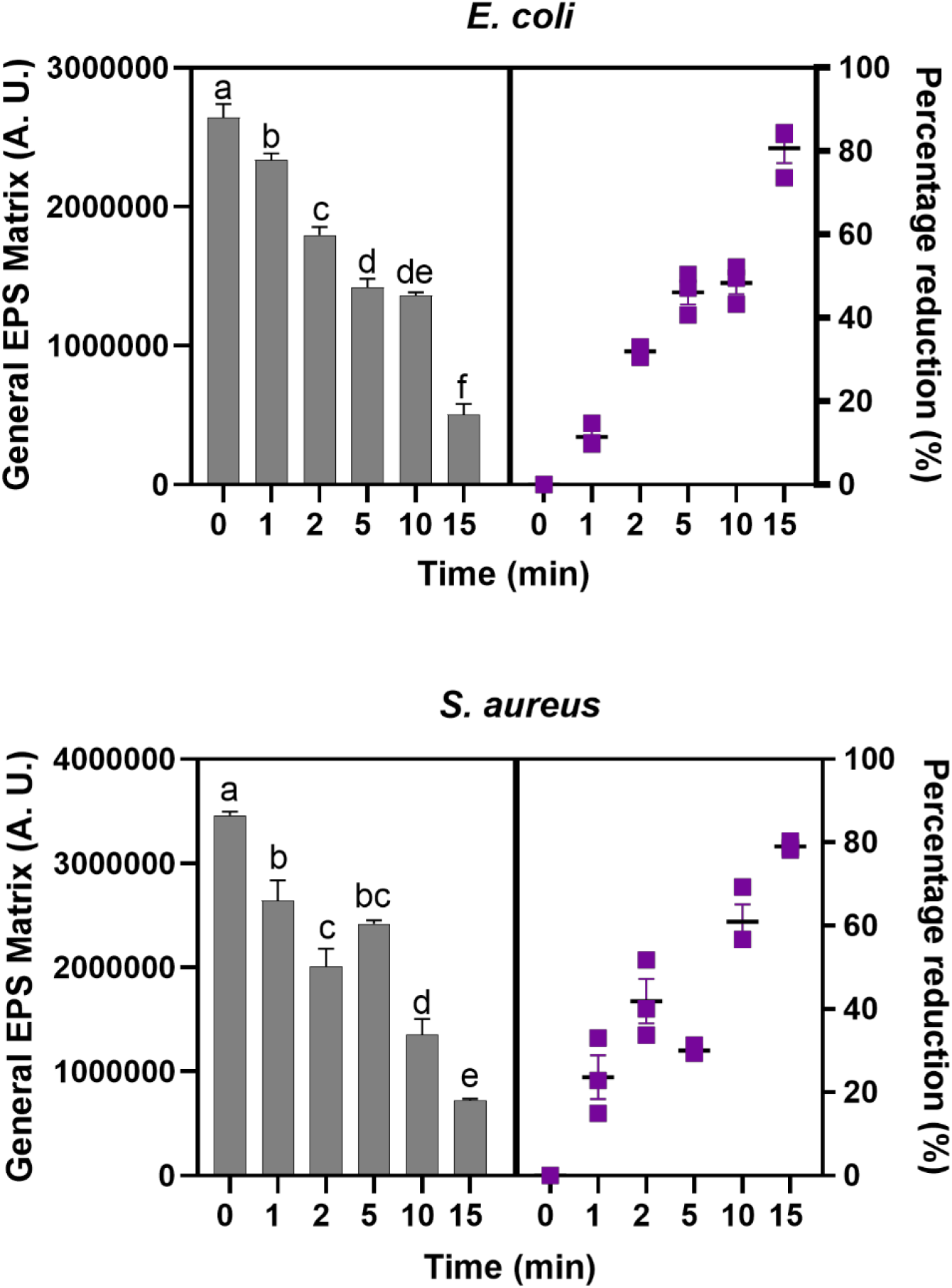
Time-dependant general EPS matrix deduction of *E. coli* UTI89 and *S. aureus* NCTC8325 biofilms challenged by 0 to 15 min of *in situ* PAW treatment. Different letters indicate significant differences between groups (P<0.05). Data represents mean ± SEM; n = 3.

In addition to the quantitative polysaccharides assessment, the PAW treated biofilms were stained with fluorescent probes to visualize for polysaccharides, eDNA and protein components of the EPS matrix using a fluorescent microscope (Figure 5). The polysaccharides (Con A staining) of *E. coli* and *S. aureus* biofilms exhibited a significant decrease compared to the untreated biofilms after 1, 5, and 15 min treatment. From the images, some polysaccharide debris was observed after PAW exposure, confirming the results of the quantitative EPS matrix evaluation above.

**Figure 5.**
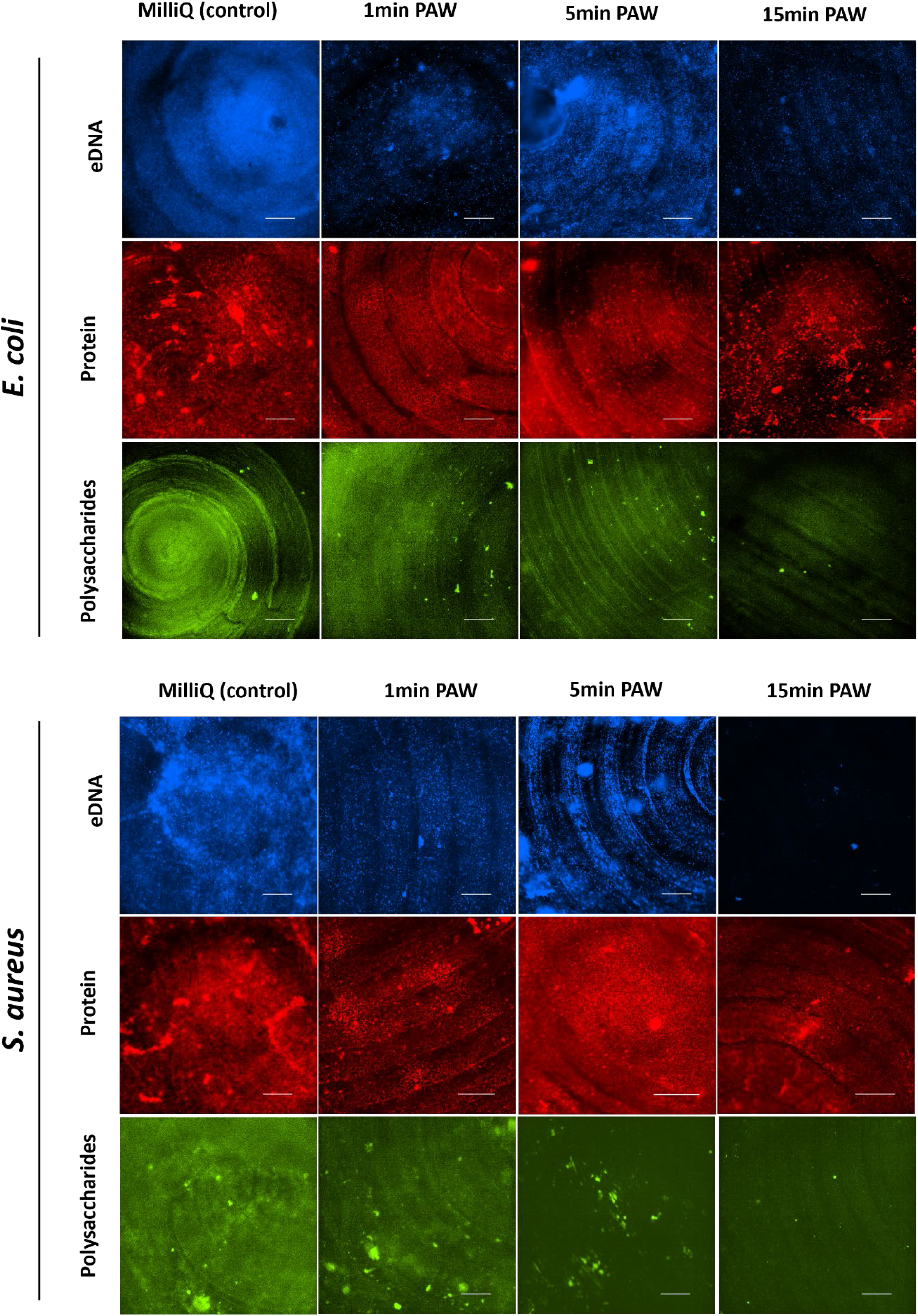
The EPS matrix fluorescent images of *E. coli* UTI89 and *S. aureus* NCTC8325 under non-treatment (control), 1 min, 5 min, and 15 min-PAW treatment. EPS matrix of extracellular DNA, protein, and polysaccharides were stained using Sytox Blue, Sypro Ruby, and Con A Alexa fluo 647, respectively. Experiments were repeated in triplicates and random positions were taken. Scale bar = 100 µm.

Sytox Blue, impermeable to live cells but able to stain the extracellular nucleic acid of dead cells or the matrix, was utilized for staining eDNA. The fluorescence intensity of the eDNA stain in *E. coli* and *S. aureus* decreased rapidly after 1 min treatment, followed by a notable increase in fluorescence intensity at 5 min treatment, after which its fluorescent signal significantly decreased. The observed increase in blue fluorescence at 5 min treatment might be attributed to the DNA released by dead cells following PAW exposure.

In contrast to the consistent reduction observed in exopolysaccharides and eDNA, the protein matrix in *E. coli* and *S. aureus* biofilms remained present even when cells were undetected. As shown in the images under red fluorescence in Figure 5, extracellular proteins were clustered and evenly spread across the steel surface. Following 1 min treatment, a significant portion of these protein clusters was removed, revealing an evenly distributed protein signal resembling the shape of cells. Subsequently, an increase in protein fluorescence was noted at the 5 min-PAW treatment point.

### 3.4 Biofilm morphology changed after PAW treatment

To visualise the effects of *in situ* PAW treatment on the biofilm and the cell structure, SEM imaging was conducted (Figure 6). As biofilm cells were completely removed after 15 min treatment, therefore, cells under 1 min and 5 min PAW treatment were imaged by SEM. Qualitative analysis demonstrates a discernible decrease in the number of physically adhered *E. coli* and *S. aureus* biofilm cells on the stainless-steel surface following both 1 min and 5 min PAW treatments, as compared to the MilliQ control. Additionally, PAW treatments induce morphological alterations in both *E. coli* and *S. aureus* biofilm cells, resulting in a flattened appearance that is particularly evident in *E. coli* cells.

**Figure 6.**
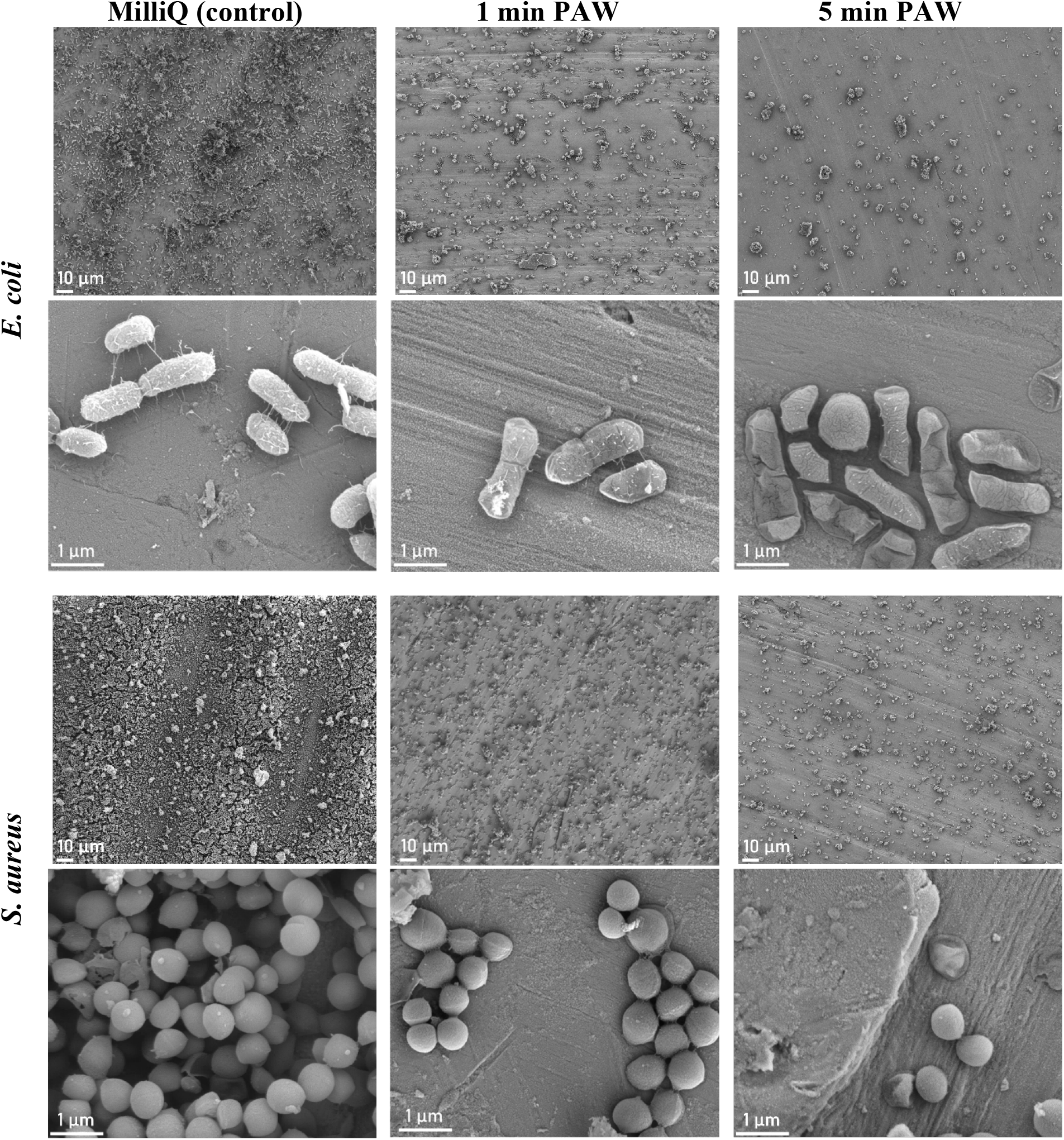
PAW treatment physically dislodges *E. coli* and *S. aureus* bacterial biofilm cells and is altering the bacterial cell morphology. SEM imaging was utilised to visualise the architectural and morphological changes induced by 1 and 5 min treatments. A MilliQ control was included for comparison. Both 1 and 5 min treatment time points resulted in significant bacterial cell removal from the stainless steel surface, with some cells exhibiting a flattened cell morphology.

### 3.5 Membrane response to PAW

The alterations in membrane permeability and extracellular membrane potential enable bacteria to respond to environmental stress by connecting intracellular reactions to external conditions (Mingeot-Leclercq and Décout 2016). Following this principle, the NPN-based assay and DiSC_3_(5) stains were used to elucidate the mode of action of PAW on the membrane permeability and membrane depolarization of *E. coli* and *S. aureus* cells (Figure 7). NPN fluorescence increases as the dye enters the cell due to the disruption of the membrane that leads to an increase in permeability. In PAW-treated *E. coli* and *S. aureus* suspensions a time-dependent increase in the fluorescence of NPN was observed. Compared to the MilliQ control, the fluorescence of NPN of *E. coli* and *S. aureus* suspensions rapidly increased after approximately 1 min treatment. NPN uptake fluctuated but remained not significantly different until 30 min.

**Figure 7.**
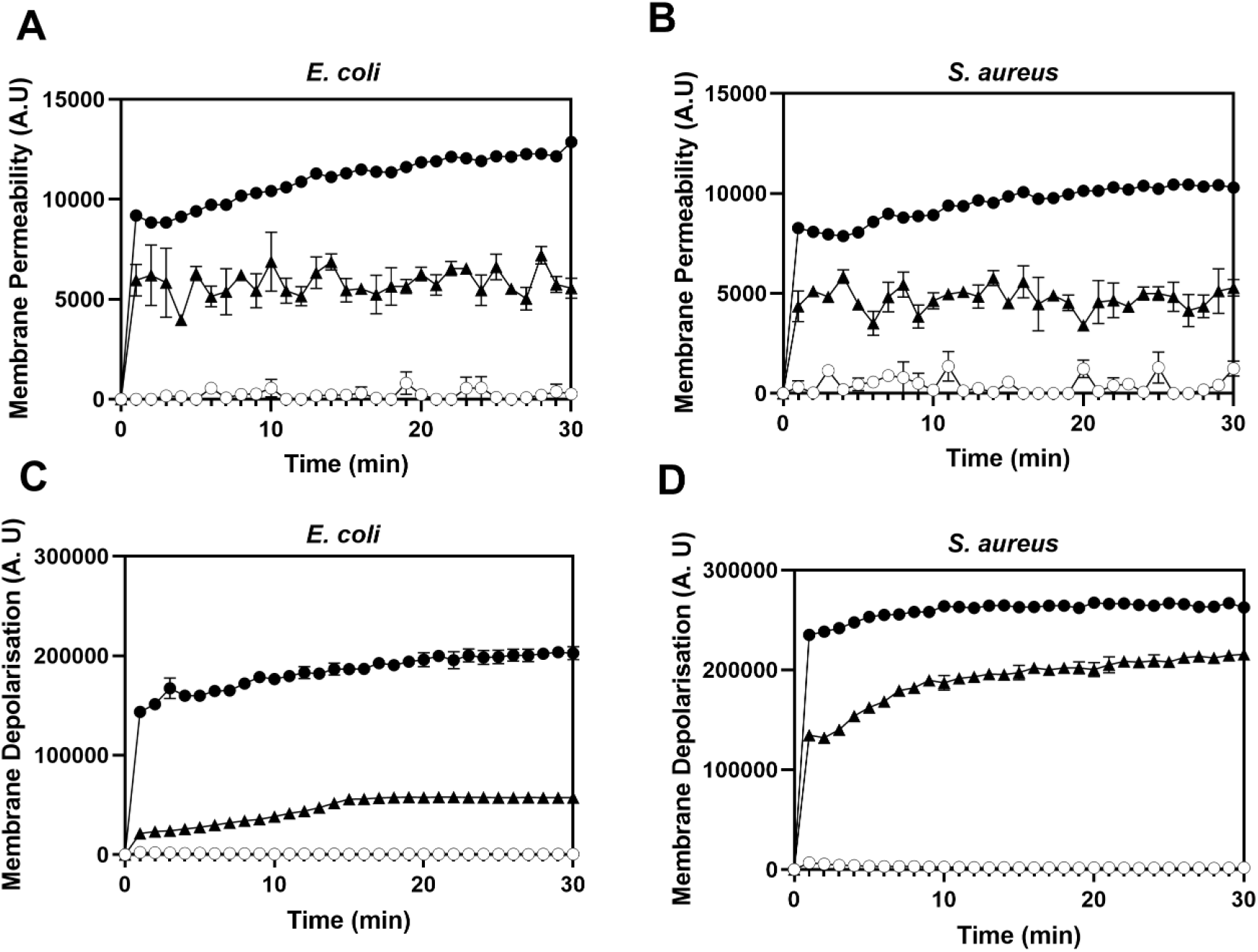
The response of bacterial *E. coli* UTI89 and *S. aureus* NCTC 8325 membrane permeability (A and B: assayed by NPN staining) and membrane depolarisation (C and D: assayed by DiSC3(5) staining) in the presence of control (negative control of Milli-Q water(○), PAW (▴), and CTAB (positive control,(•).

DiSC_3_(5) can evaluate the polarization state of cells, as it can penetrate the lipid bilayer of membrane and further accumulate in cells intracellularly (Buttress 2022). An increased fluorescence of DiSC_3_(5) under PAW treatment was observed in both of *E. coli* and *S. aureus* cells (Figure 7). More specifically, the membrane of *S. aureus* cells exhibited significant depolarization at 1 min PAW exposure (P≤0.0001). This effect gradually increased until 20 min and remained consistent until 30 min. Conversely, PAW induced a less pronounced depolarization of the *E. coli* cell membrane compared to *S. aureus*. The membrane of *E. coli* gradually depolarized until 15 min (P<0.05), with this effect remaining stably until 30 min compared to the MilliQ control.

### 3.6 PAW induced an increasing of intracellular RONS level

The accumulation of intracellular ROS and RNS within the PAW-treated *E. coli* and *S. aureus* biofilms was assessed using fluorescence probes DCFDA (ROS, Figure 8A) and DAF-FM (RNS, Figure 8B), respectively., A significant increase in DCFDA fluorescence (P ≤ 0.0001) was observed in both *E. coli* and *S. aureus* biofilm cells compared to the control, indicating that 15 min-PAW treatment induced the accumulation of ROS within both *E. coli* and *S. aureus* biofilms. However, the amount of intracellular ROS increased even more significantly (P ≤ 0.01) in *S. aureus* biofilms compared to *E. coli* biofilms. The fluorescence of DAF-FM-stained *E. coli* and *S. aureus* biofilms challenged by PAW treatment was also significantly increased (P ≤ 0.0001), but there was no significant difference in intracellular RNS accumulation between *E. coli* and *S. aureus* biofilm.

**Figure 8.**
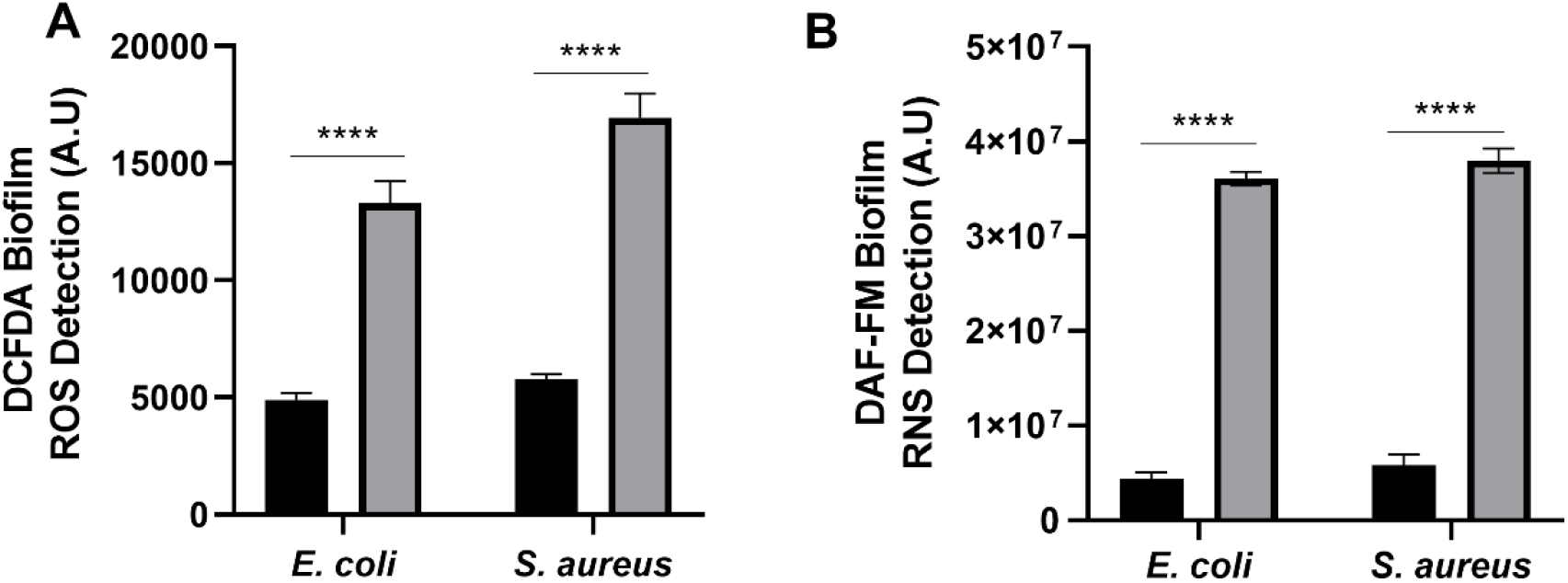
PAW induces the intracellular accumulation of ROS and RNS within bacterial (*E. coli* UTI89 and *S. aureus* NCTC8325) biofilm cells. (A) DCFDA staining assay and (B) DAF-FM staining assay was performed to detect the intracellular ROS and RNS under 15 min-PAW treatment (grey bar). Milli Q water was applied as the control (black bar). Data represents mean ± SEM, * (P<0.05), **(P<0.01), ***(P<0.001), and ****(P<0.0001); n =3 biological replicates, with 3 technical replicates for each.

## 4 Discussion

### 4.1 The viability of biofilm cells and biofilm biomass was significantly reduced by *in-situ* PAW treatment

The antibacterial efficacy of PAW varies among different bacterial strains. In this study, PAW was able to eradicate biofilm cells completely for both of *E. coli* and *S. aureus*. However, this process happened 2 min quicker in *S. aureus* than *E. coli*. Previous studies have indicated that Gram-negative bacteria (such as *E. coli* and *P. aeruginosa*) are generally less resistant to PAW treatments compared to Gram-positive bacteria (such as *Bacillus subtilis* and *Staphylococcus. epidermidis*) (Hozák 2018; Mai-Prochnow 2016; Ziuzina 2015). However, non-selective activity of PAW on both *S. aure*us (Gram +) and *P. aeruginosa* (Gram -) has also been reported (Brun 2018). In contrast to our findings, it has been observed that gram-positive cells are more resistant to gaseous plasma treatment, likely due to the physical protection provided by the thicker peptidoglycan layer of gram-positive cells (Mai-Prochnow 2021). The variations in antibacterial biofilm efficacy reported may be attributed to differences in the chemical reactive species present in PAW generated under different systems and the structural differences among bacterial cells.

Despite the absence of viable CFU cells of *E. coli* and *S. aureus* biofilms after 15 min-PAW treatment (Figure 2), approximately 50% of the biofilm biomass remained on the coupon surface. This suggests that while PAW treatment kills all biofilm cells, it may only remove a portion of the bacterial biofilm structure from the steel surface. Several studies have similarly found that the biofilm structure persists, and biofilm biomass retained the same or even slightly increased after PAW treatment (Chen 2017; Hozák 2018). This phenomenon might be due to increased metabolic activities caused by a high concentration of NO_3_^-^ in PAW used as nitrogen nutrition (Chen 2017). Together, it could be hypothesized that the reactive stress from PAW may detach a portion of the biofilm biomass, exposing the biofilm cells, and allowing diffusion of reactive plasma species into biofilms through the protective matrix that then leads to the killing of the inner cells. However, the mechanisms of action of PAW on the bacterial biofilm EPS matrix remain unclear and warrant further investigation.

### 4.2 Antimicrobial mechanisms of PAW against bacterial biofilm

The impact of PAW on the EPS matrix exhibits variability across different components. In this study, we observed a notable removal of eDNA and polysaccharides from the EPS matrix. eDNA serves as a cohesive agent, binding the biofilm matrix with the cells, which is crucial for structural stability (Flemming and Wingender 2010). Upon eDNA removal, the biofilm cells are more prone to detachment from the matrix, rendering them more susceptible to oxidative exposure from PAW (Patange 2021). The observed changes in eDNA levels suggest that PAW initiates biofilm cell detachment as early as the first minute of treatment by removing eDNA, subsequently disrupting the bacterial cells to release intracellular DNA, thus facilitating the complete eradication of biofilm cells.

We observed an increase of eDNA matrix at 5 min-PAW treatment on both of *E. coli* and *S. aureus* biofilm matrix. One possible explanation could be the release of proteins by dead cells, akin to the eDNA dynamics mentioned earlier, as cell wall comprises a significant proportion of glycoproteins. Oxidative agents in PAW (e.g., hydrogen peroxide, a long-lived ROS generated in PAW) has demonstrated the ability to oxidating proteins, disrupting protein and DNA (da Cruz Nizer 2021). Additionally, the reactive species in PAW might inhibit the enzyme activity of producing catalase enzymes and superoxide dismutase which can convert harmful ROS within the cell before they induce damage (Finnegan 2010).

The significant removal of extracellular polysaccharides from the EPS matrix might contribute to the oxidative reaction between PAW RONS to the cells within biofilm, leading to their destruction. For instances, the hydroxyl and carboxyl groups in polysaccharides may trap the plasma-generated OH radicals, resulting in hydrogen abstraction and subsequent molecular damage to polysaccharides (Khosravian 2014). This partially explains the reduction in biofilm biomass through PAW treatment, although the question regarding the remaining biomass with undetectable biofilm cells remains unresolved.

PAW treatment involves physical dislodgement as well as flattening of the individual cells. These physical effects are likely due to the reactive oxygen species present in the PAW, with ROS known to disrupt the microbial membrane (Heema Kumari Nilesh 2019; Rothwell). We have previously shown that within 1 min-PAW treatment, *E. coli* biofilm cell membranes are rapidly depolarised, and the outer membrane is significantly permeabilised (Vyas 2023). Other research also found that the outer membrane of *S. aureus* cells was significantly damaged due to the oxidative stress created in cells during PAW treatment (Rathore 2021). However, as noted by Vyas et al., (2023) PAW that was applied post-generation did not physically remove biofilm, suggesting that the *in situ* treatment utilised in this study has the additional benefits of physical forces that can weaken and disrupt biofilm structures. This highlights an advantage of *in situ* treatment, as it may enhance biofilm susceptibility to subsequently applied antimicrobials (e.g., antiseptics, disinfectants).

The oxidative stress (ROS and RNS) created in cells during PAW results in cell death (Gunes 2022). Plasma-induced membrane damage involves the cumulative impact of several long- and short-lived ROS, such as superoxide anion radicals, hydroxyl radicals, hydrogen peroxide, and ozone. These can act on membrane-associated proteins, further triggering oxidative stress within biofilm cells, and DNA damage within cells (Balcerczyk 2005; Carpenter and Schoenfisch 2012). We have previously shown that PAW generated with air (mixture of mostly nitrogen and oxygen, some argon) as an input gas source results in significant intracellular RONS accumulation for E. coli biofilms (Vyas 2023), whilst ROS significantly predominate within biofilms treated with PAW that is generated with pure oxygen (Xia 2023). Further studies should examine PAW generated with differing input gas sources (e.g., pure oxygen, nitrogen or argon) for their effects on intracellular RONS accumulation and how biofilm viability, biomass, and EPS may be impacted differently.

## 5 Conclusion

This study demonstrated that PAW treatment exhibits a generally non-selective inhibitory effect on pathogenic *S. aureus* NCTC8325 biofilms and *E. coli* UTI89 biofilms. The study also contributes novel insights into PAW’s mechanisms of action, particularly its impact on the EPS matrix (exopolysaccharides, extracellular DNA, and protein), membrane permeability, depolarization, and intracellular ROS and RNS accumulation in both Gram-positive and Gram-negative species. These findings highlight PAW’s potential as a promising strategy for biofilm control across diverse microbial species. Further research into the specific interactions between PAW components and the EPS matrix will provide valuable insights for optimizing PAW-based treatments against biofilm-related challenges in antimicrobial development and water system decontamination.

## 6 Acknowledgements

We thank Prof Dee Carter (University of Sydney) for kindly provide the *S. aureus NCTC8325* stains utilized in this study and thank Dr. Parisa Doorian (University of Technology Sydney) for kindly provide the *E. coli UTI89* stains utilized in this study. We thank Nicolas Gracie (University of Sydney) for their invaluable support of providing the Nikon fluorescence microscopy equipment. We thank Sydney Microscopy & Microanalysis (University of Sydney) for accessing the Scanning electron microscope.

## 7 Funding

This work was supported by the Australian Research Council Discovery Scheme (DP210101358).

## 8 CrediT

Binbin Xia: Conceptualization, Methodology, Investigation, Validation, Data curation, Formal analysis, Writing – original draft. Heema Kumari Nilesh Vyas: Investigation, Methodology, Writing – original draft, review & editing. Scott A. Rice: Writing – review & editing, Project administration, Funding acquisition. Timothy P. Newsome: Writing – review & editing. Patrick J. Cullen: Supervision. Anne Mai-Prochnow: Conceptualization, Supervision, Writing – review & editing, Funding acquisition, Project administration.

## 9 Conflict of Interest

The authors declare the following financial interests/personal relationships which may be considered as potential competing interests: Author PJ Cullen is the CTO of PlasmaLeap Technologies, the supplier of the plasma power source and reactors employed in this study.

